# Numerical investigation of heterogeneous soft particle pairs in inertial microfluidics

**DOI:** 10.1101/2023.08.24.554596

**Authors:** Benjamin Owen, Krishnaveni Thota, Timm Krüger

**Affiliations:** School of Engineering, Institute for Multiscale Thermofluids, University of Edinburgh, Edinburgh EH9 3FB, UK

## Abstract

The formation of pairs of particles or cells of different types in microfluidic channels can be desired or detrimental in healthcare applications. It is still unclear what role softness heterogeneity plays in the formation of these particle pairs. We use an in-house lattice-Boltzmann-immersed-boundary-finite-element solver to simulate a pair of particles with different softness flowing through a straight channel with a rectangular cross-section under initial conditions representative of a dilute suspension. We find that softness heterogeneity significantly affects the pair dynamics, determining whether a pair will form or not, and determining the lateral and inter-particle equilibrium behaviour in the pair. We also observe close matches between the transient deformation of particles in a linear pair and single particles in isolation. These results further our understanding of pair behaviour, providing a foundation for understanding particle train formation, and open up the potential to develop reduced-order models for particle pair formation based upon the behaviour of single particles.

## 1 Introduction

Pathological alterations in cell properties are associated with various diseases, such as malaria ^1^ and sickle cell anaemia ^2^. These alterations in cell properties can be exploited in disease diagnostics ^3^ and therapeutic tools ^4^. Inertial microfluidic (IMF) devices are able to passively manipulate and separate cells based on their properties, such as size, shape, and softness, by focusing cells into axially ordered trains. IMF devices offer advantages over traditional microfluidic methods due to higher throughput, lower cost, and the ability to manipulate particles in a label-free manner ^5^.

Inertial microfluidics was first proposed by Di Carlo *et al*. in the late 2000s ^6,7^. The basic premise is to increase the fluid inertia within the microfluidic device (channel Reynolds number of order 10–100), usually by increasing the flow rate. In doing so, not only does throughput of cells under analysis increase, but also inertial effects can be exploited to manipulate particles through focusing and separation ^8,9^.

The particle focusing phenomenon was first observed by Segré and Silberberg in a pressure-driven flow through a cylindrical pipe ^10^. A single particle in a straight channel will focus (migrate laterally) to a cross-sectional equilibrium position. This focusing is caused by the balance of two main forces, a shear gradient lift force, and a wall-induced repulsion force. The shear gradient lift force usually pushes the particle away from the channel centre toward the walls. Contrarily, the wall-induced repulsion force pushes the particle toward the centre of the channel. The balance of both forces dictates the location of the lateral equilibrium positions.

Where more than one particle exists in an inertial microfluidic device, these particles interact hydrodynamically. For example, the existence of other particles causes changes in the flow field which can modify the shear gradient lift force and therefore modify the lateral equilibrium positions. Hydrodynamic interactions in the inertial regime also cause an additional phenomenon, the axial ordering of particles into trains with regular inter-particle spacing ^11,12^. These trains can be exploited in applications such as cell encapsulation ^13^ and flow cytometry ^14^.

The formation mechanism of particle trains is not fully understood. However, Schaaf *et al*. ^15^ demonstrated that particles form distinct pairs that later join together to form trains. It is therefore crucial to understand the formation mechanism of particle pairs.

Particle pairs can be classified into staggered pairs (particles located on opposite sides of the channel) and linear pairs (particles located on the same side of the channel) ^15,16^. The self assembly of particle pairs due to reverse streamlines was first identified by Lee *et al*. ^16^. During pair formation, the axial distance between both particles converges to an equilibrium value ^11,17^. It has been reported that linear particle pairs do not form when both particles are of the same size ^15,18^. In previous work ^19^, we reported that particles form homogeneous pairs from a larger range of initial positions when the particles are softer.

While we know how softness affects pair formation when both particles have the same softness, the effect of softness heterogeneity, as it would be expected in real-world applications, is less clear. Patel and Stark ^20^ investigated the effect of particle softness for mono- and bi-disperse particle pairs and found that the presence of the second particle can change the stability of the single-particle equilibrium positions. Li *et al*. ^21^ investigated the formation of a heterogeneous particle pair consisting of a rigid and a soft particle and demonstrated the pair formation after a number of passing interactions in a simulation with periodic boundary conditions. However, in real-world applications, passing particles would not interact repeatedly, thus, it is important to study the outcome of the first interaction between two particles.

The formation of heterogeneous pairs and trains can be desired (for example, for the generation of compound particles) or detrimental (for example, for the separation of different particle species). Thus, it is important to better understand the conditions and mechanisms leading to the formation of pairs of particles with different softness. In this paper, we perform 3D simulations using a lattice-Boltzmann-immersed-boundary-finite-element solver to numerically investigate the dynamics and formation of a pair of particles of different softness flowing through a straight rectangular channel at a moderate Reynolds number (Sec. 2). We consider particle pairs in the staggered and linear arrangements (Sec. 3). Our study finds that softness heterogeneity strongly affects the pair dynamics. We show that linear pairs can form when particles are different, and we demonstrate that particles deform more during close particle-particle interactions than when in isolation. We demonstrate that particle deformation during the formation of linear pairs can be predicted from the single-particle behaviour, while the deformation during the formation of staggered pairs is more complicated. Implications and future directions are discussed in Sec. 4.

## 2 Methods and system characterisation

The physical and numerical models are briefly outlined in Sec. 2.1 and Sec. 2.2, respectively. Sec. 2.3 describes the analysis of paticle deformation, and Sec. 2.4 summarises the typical particle trajectories observed.

### 2.1 Physical model

#### 2.1.1 Microfluidic set-up and governing equations

We consider a Newtonian liquid with kinematic viscosity *ν* and density *ρ* flowing through a straight channel with a rectangular cross-section with width 2*w* and height 2*h* as shown in Fig. 1. The flow is periodic along the flow direction (*x*-axis), and the length of the periodic unit cell is *L*. The fluid flow is pressure-driven and governed by the incompressible Navier-Stokes equations. The *y*- and *z*-axes are denoted as lateral directions.

**Fig. 1.**
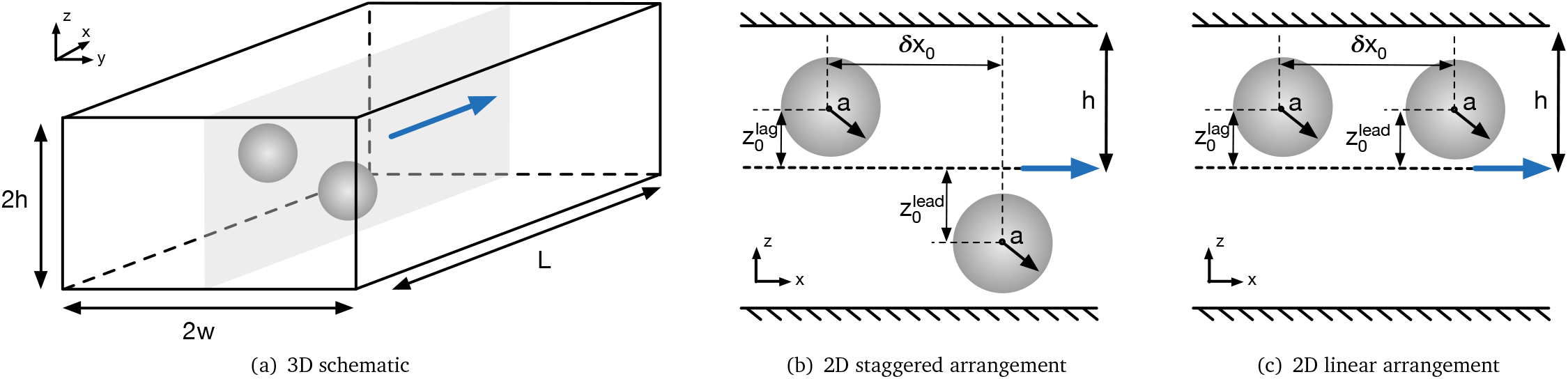
Schematic of particle pairs in a straight, rectangular duct with height 2*h* and width 2*w*. The periodic unit cell length is *L*. The flow is along the *x*-axis (blue arrow). Particles are initially located on the mid-plane with *y* = const (indicated by grey plane). (b) Particles are shown in a staggered arrangement where particles are located on different sides of the channel. (c) Particles are shown in a linear arrangement where particles are located on the same side of the channel.

We consider two spherical, neutrally buoyant particles with radius *a*. The suspended particles are modelled as deformable capsules comprising a thin hyperelastic membrane and interior liquid with the same properties as the suspending liquid. We employ the Skalak model for the capsule membranes ^22^:

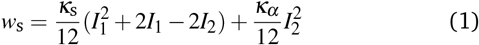

where *w*_s_ is the areal energy density, *I*_1_ and *I*_2_ are the in-plane strain invariants ^23^, and *κ*_s_ and *κ*_*α*_ are the elastic shear and area dilation moduli. We include a membrane bending energy

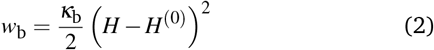

where *H* and *H*^(0)^ are the trace of the surface curvature tensor and the spontaneous curvature, respectively, and *κ*_b_ is the bending modulus.

Both particles are initialised on the mid-plane of the longest cross-sectional axis (*y* = const) as shown in Fig. 1a, while the initial *x*- and *z*-coordinates are varied. Particles initially located on the mid-plane will remain on this plane while moving along the *x*-axis and migrating along the *z*-axis ^24^. We have not observed particles leaving the mid-plane in any of our simulations. We consider two particle arrangements in this work. In the staggered arrangement, the particles are placed on opposite sides of the channel (Fig. 1b). In the linear arrangement, the particles are placed on the same side of the channel (Fig. 1c). In both cases, We distinguish between the initially leading (farther downstream) and lagging (farther upstream) particles, according to their initial positions along the flow axis (*x*-axis).

We apply periodic boundary conditions in the axial direction and the no-slip boundary condition at the channel wall and the surfaces of the particles.

#### 2.1.2 Characteristic scales and dimensionless groups

The channel Reynolds number is defined as

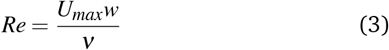

where *U*_*max*_ is the maximum fluid velocity in the undisturbed flow (flow without particles).

The Laplace number is used to characterise the softness of the particle and is defined as the ratio between the typical elastic shear forces in the capsule membrane and the intrinsic fluid force:

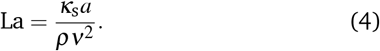

As the Laplace number is a combination of material properties only, it is suitable to isolate the contribution of particle softness in inertial flows ^25^.

Other dimensionless groups are the channel aspect ratio *w/h*, particle confinement *χ* = *a/h*, the reduced dilation modulus 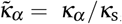, and the reduced bending modulus 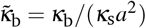.

We use particle radius *a*, or channel half-height *h* to non-dimensionalise distances and positions. Time is non-dimensionalised by the advection time of the particle:

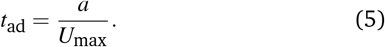

In this study we keep the following dimensionless groups constant: the channel Reynolds number (*Re* = 10), channel aspect ratio (*w/h* = 2), and particle confinement (*χ* = 0.4).

### 2.2 Numerical model

The numerical model consists of a partitioned fluid-structure interaction solver in which the lattice Boltzmann (LB) method is used for the fluid flow, the finite element (FE) method for the capsule dynamics, and the immersed boundary (IB) method for the fluid-structure interaction. This IB-LB-FE solver has previously been employed in the study of the dynamics of deformable red blood cells and capsules ^26,27^. Here, we provide essential properties of the model, while comprehensive details are available elsewhere ^23^.

For the LB method, we use the D3Q19 lattice ^28^ and the BGK collision operator ^29^ with relaxation time *τ*. The kinematic viscosity of the liquid and the relaxation time satisfy

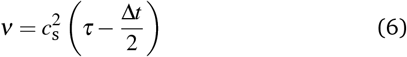

where *c*_s_ is the lattice speed of sound and Δ*t* is the time step. For the D3Q19 lattice, 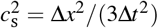 holds where Δ*x* is the lattice resolution. For the channel flow cases, flow is driven by a constant body force following the forcing method of Guo *et al*. ^30^. This form of the LB method is widely used in the field of fluid dynamics, including in previous inertial microfluidics studies ^25,31^.

Each capsule is represented by a surface mesh with *N*_f_ flat triangular faces (or elements) defined by three nodes (or vertices) each. At a given time step, the capsule mesh is generally deformed. The hyperelastic forces acting on each vertex are calculated as a function of the mesh deformation state through an explicit scheme. The shear and area dilation forces result from the deformation gradient tensor of each face, while the bending forces are related to the angles between normal vectors of pairs of neighbouring faces ^26^.

We employ an IB method with a 3-point stencil ^32^. The forces obtained from the FE scheme are spread from the Lagrangian mesh to the Eulerian lattice where they act on the surrounding fluid nodes through the LB algorithm. The updated fluid velocity is then interpolated at the location of each mesh node. The positions of the mesh nodes are updated using the forward-Euler method, assuming a massless membrane which is appropriate for neutrally buoyant capsules. This treatment recovers the no-slip boundary condition at the surface of the capsules and the momentum exchange between the liquid and the capsule membrane.

The no-slip boundary condition at the resting and moving walls for channel and shear flow, respectively, is realised by the standard half-way bounce-back condition ^33^. The flow is periodic along the *x*-axis. The channel length *L* is chosen sufficiently long to avoid the interaction of capsules with their periodic images. In all simulations, the hydrodynamic forces are sufficient to prevent contact between capsules such that capsules do not come closer than about 2Δ*x* and a repulsion force between capsules is not required. Likewise, capsules always keep a large distance from the confining walls due to hydrodynamic lift, and an additional capsule-wall repulsion force is not needed.

### 2.3 Characterisation of particle deformation

Soft particles are deformed by the surrounding flow field which is determined by both the background flow and the flow distortions caused by other nearby particles. We use the Taylor deformation index, *D*, to characterise the deformation state:

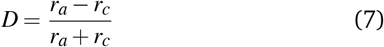

where *r*_*a*_ and *r*_*c*_ are the maximum and minimum semi-axes of the ellipsoid with the same tensor of inertia as the deformed particle ^34^. In this study, we investigate soft particles that are spherical in their undeformed state, corresponding to *D* = 0.

### 2.4 Trajectory types

We provide a brief overview of the trajectory types observed in our previous study. Six different types of trajectories have been identified in homogeneous pairs of soft particles: *Capture, Scatter, Swap & Capture, Swap & Scatter, Pass & Capture* and *Pass & Scatter*. A full description of each trajectory type can be found in Owen and Krüger ^19^. These trajectories can be categorised based on two characteristics. The first characteristic is whether the particles are captured (bound together hydrodynamically) or scattered. This characteristic is important since a stable pair cannot form unless the particles are captured. Note that not all captured particles form a stable pair since some pairs have oscillating or fluctuating distances within certain bounds. The second characteristic is whether a close particle-particle interaction occurs (either one particle passing the other or the exchange of lateral positions as one particle pushes the other — *Swapping*). This characteristic is important since a particle-particle interaction can perturb the system, moving particles away from their equilibrium positions. In suspensions with more particles, perturbations caused by additional particles could cascade and cause further interactions with other particles.

## 3 Results and Discussion

We analyse the interaction of a pair of particles with different softness, characterised by the Laplace number. Particle interactions can end in scattering (Sec. 3.1) or capturing (Sec. 3.2). The transition between both regimes is more closely investigated in Sec. 3.3.

### 3.1 Pair scatter

We begin our study by investigating the behaviour of two particles with different softness, La = 10 and La = 100. Both particles are initially located at their respective lateral equilibrium positions (also denoted as single-particle equilibrium positions), 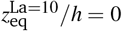 and 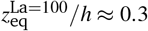. Single-particle equilibrium positions were obtained through separate simulations involving a single particle in the same geometry and under the same conditions as in Tab. 1 (data not shown). Note that, since one particle is at the centreline in this particular case, there is no distinction between the linear and staggered arrangements. The initial axial inter-particle spacing is *δx*_0_ = 10*a* and sufficiently large so that particles do not interact initially ^18,19^. These initial conditions are representative of likely scenarios within a dilute suspension where particles are rarely close to each other and have had time to equilibrate before approaching each other. We arrange the particles in both possible orders: *Leading Soft*, where the leading particle is softer, and *Leading Stiff*, where the leading particle is stiffer.

The time evolution of the lateral position and axial interparticle spacing are shown in Fig. 2(a) and (b) for both the *Leading Soft* (grey) and *Leading Stiff* (blue) configurations, respectively. In neither configuration a stable pair forms.

**Fig. 2.**
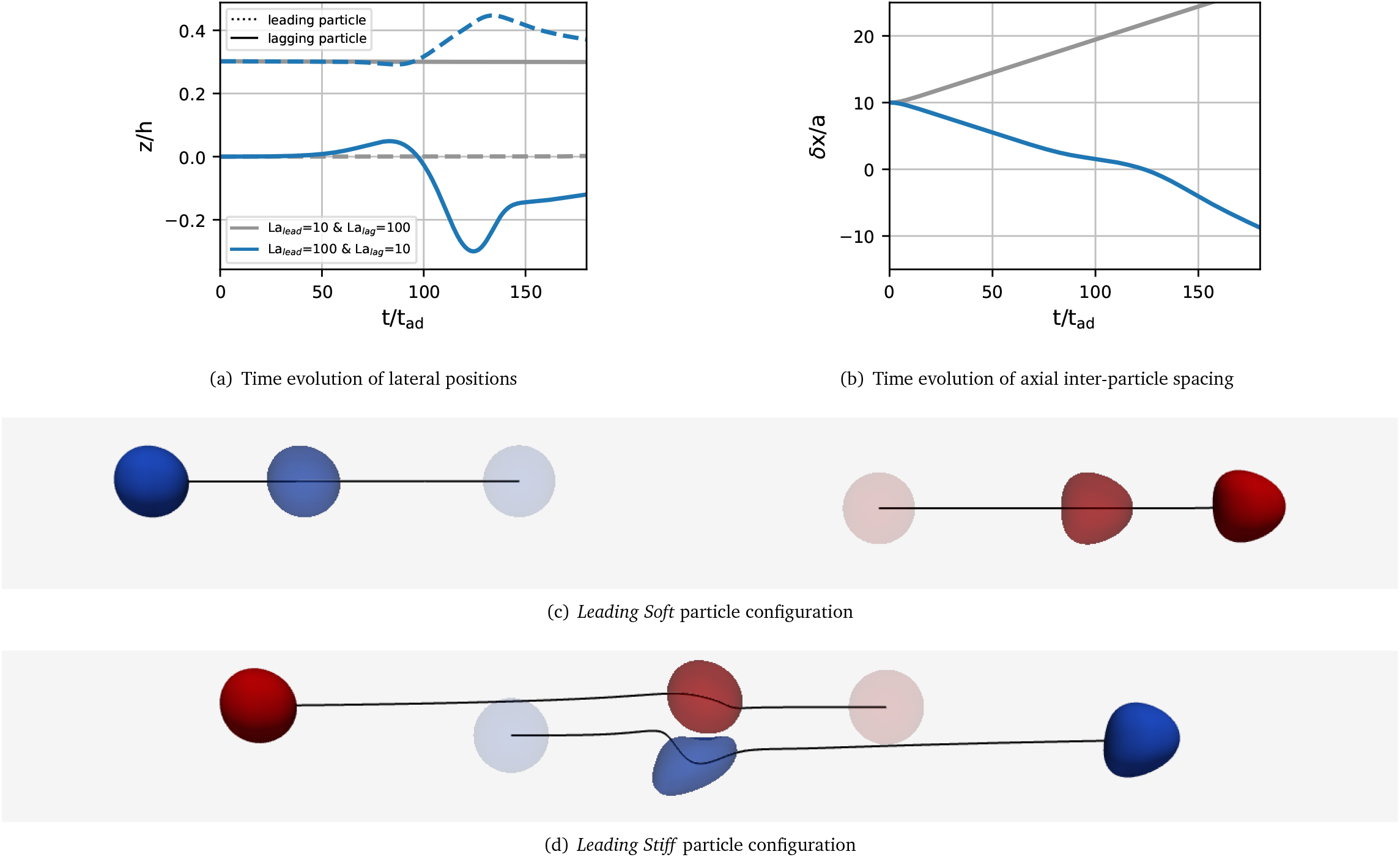
Time evolution of the (a) lateral positions of the leading and lagging particles and (b) axial inter-particle spacing for the *Leading Soft* (grey) and *Leading Stiff* (blue) configurations for a pair with La = 10 and 100. Particles are initially placed at their respective lateral single-particle equilibrium positions, and the initial axial inter-particle spacing is 10*a*. Note that, since the softer particle is initially at the centreline, there is no distinction between the linear and the staggered arrangements. Visualisations of particle configurations in (c) the *Leading Soft* and (d) the Leading Stiff configurations in the centre-of-mass frame. Particles are coloured with respect to their initial positions: the leading particle is red, and the lagging particle is blue. Particles at earlier times are paler, and particles at later times are more saturated.

**Table 1.**
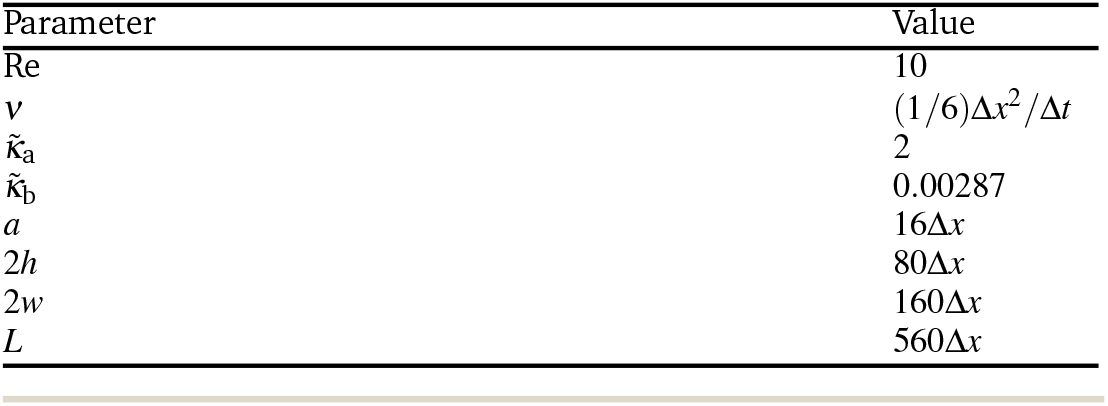
Parameters used in this study. See Fig. 1 for an illustration of the set-up. The channel Reynolds number is varied by the body force and therefore *U*_max_ via Eq. (3), and the Laplace number is controlled by the shear elasticity via Eq. (4). The liquid density is set to 1 in simulation units.

In the *Leading Soft* case, the leading particle moves away from the lagging particle and no interaction occurs. This observation is expected since the leading particle is located at the centre of the channel whereas the lagging particle is off-centre. As a result, the leading particle has a larger axial velocity and moves away.

Contrarily, in the *Leading Stiff* case, the lagging particle, being on the channel centreline, is faster than and catches up with the leading particle, which is off-centre. Upon reaching the leading particle, the lagging particle is able to push the leading particle farther from the channel centre and pass it. The initially lagging particle is now leading and moves away from the initially leading particle, without a pair forming.

Homogeneous pairs (*i*.*e*., two particles with the same La) under the same initial conditions as the heterogeneous pairs presented in Fig. 2 do not form a stable pair. Instead, since both particles have nearly the same lateral equilibrium position, they move with the same speed and roughly maintain their initial axial distance ^19^.

### 3.2 Pair capture

In order to investigate the sensitivity of the pair behaviour to the softness of the lagging particle, we perform a sweep of the softness of the lagging particle in the range La_lag_ = [10, 100]. The softness of the leading particle remains unchanged at La_lead_ = 100. The initial lateral positions of both particles correspond to their respective single-particle equilibrium positions which are shown in Fig. 3(a). For cases where La_lag_ is large enough for the lagging particle to be off-centre initially, we consider both linear and staggered pairs, *i*.*e*., 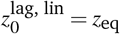 and 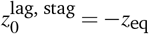. We begin with a characterisation of the spatial particle configuration and particle deformation while in a stable pair, before discussing the transient behaviour during pair formation.

**Fig. 3.**
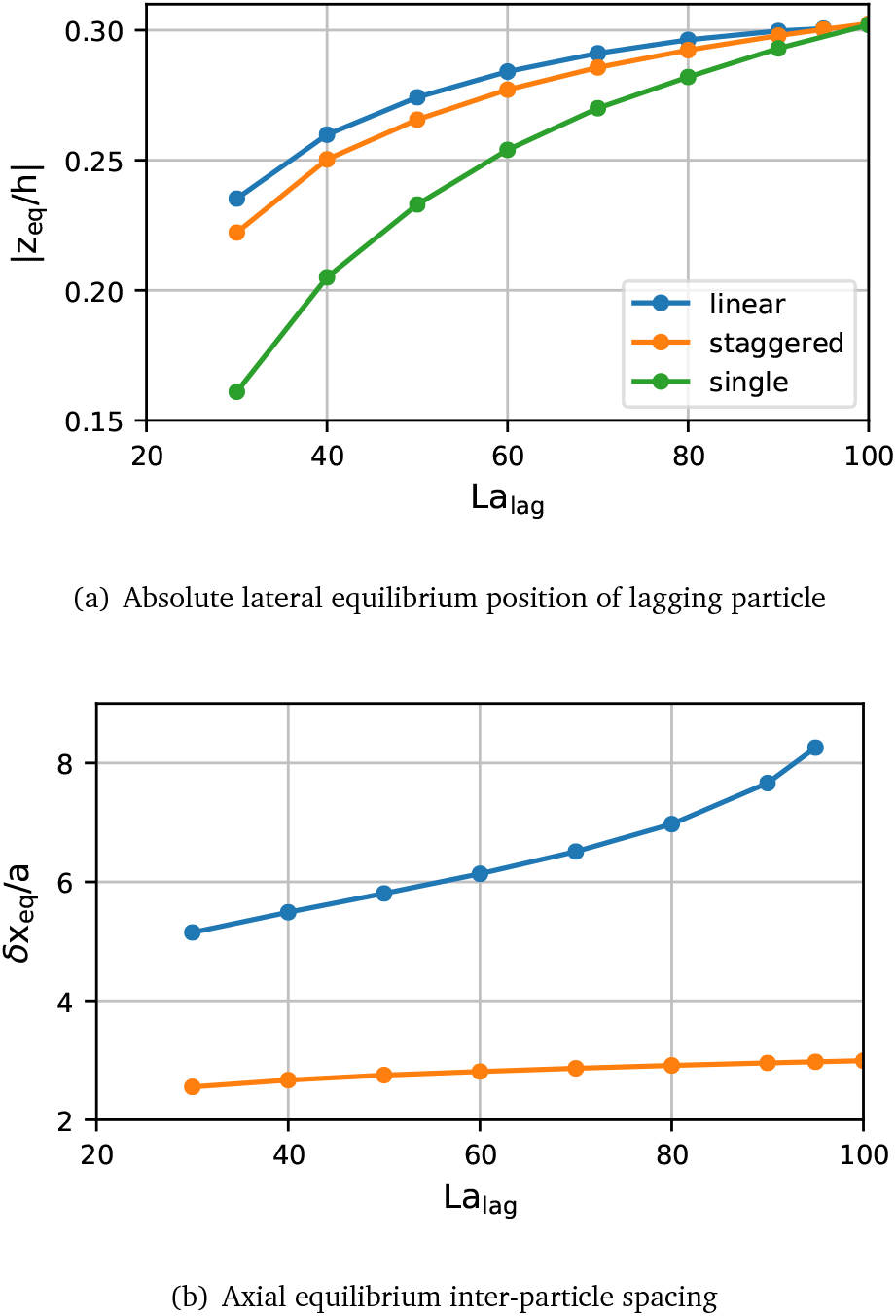
Equilibrium behaviour of stable particle pairs with La_lead_ = 100 and varying La_lag_ *∈* [30, 100]. No pairs form for smaller La_lag_. (a) Absolute lateral equilibrium position (|*z*_eq_*/h*|) of the lagging particle in a linear pair (blue), in a staggered pair (orange), and a single particle for comparison (green). (b) Axial equilibrium inter-particle spacing (*δx*_eq_*/a*) in staggered (orange) and linear (blue) pairs.

#### Spatial particle configuration in stable pairs

We find that the trajectories of pairs transition from a scatter to a capture type as the softness of the lagging particle decreases. The transition happens for La_lag_ *≈* 25, as will be discussed in more detail later. In all observed cases, the initial arrangement of the pair is maintained if a stable pair forms. Fig. 3 shows the lateral equilibrium position of the lagging particle and the axial inter-particle spacing in the pair for all stable pairs identified. A small difference in lateral equilibrium position of less than 2% exists between the leading and lagging particle in all pairs (data not shown). The difference is larger in linear pairs than in staggered pairs. We also observe that the difference decreases for both linear and staggered pairs when La_lag_ increases toward La_lead_ = 100.

As La_lag_ increases, the lateral equilibrium position of the lagging particle moves farther away from the channel centre (Fig. 3(a)). This trend is observed for both the linear and staggered arrangements. However, in all simulated heterogeneous pairs, staggered pairs converge to lateral equilibrium positions closer to the channel centre than equivalent linear pairs.

The equilibrium inter-particle spacing is more significantly affected by the pair arrangement than the lateral equilibrium position (Fig. 3(b)). The equilibrium spacing is smaller in the staggered arrangement than in the linear arrangement, confirming findings reported by Kahkeshani *et al*. ^35^. We also observe that the equilibrium spacing in both arrangements increases with La_lag_. In the staggered arrangement, this increase is nearly linear with a small variation of *≈* 0.5*a* across the investigated range of softness, while in the linear arrangement the variation is non-linear and much larger (*≈* 3.0*a*). This difference in behaviour is particularly significant in the limiting case of a homogeneous pair, La_lag_ = La_lead_: In the linear arrangement, no pair forms and the relative axial speed is very slow (*<* 0.01*a* per 1000*t*_ad_, data not shown). Conversely, a captured pair forms for particles in the staggered arrangement when La = 100 for both particles.

#### Particle deformation in stable pairs

The deformation of both particles is affected by the type of pair arrangement. Fig. 4(a) shows the Taylor deformation, *D*, in equilibrium for both particles for the same cases as in Fig. 3. The Taylor deformation for a single particle in equilibrium is included for comparison. We make several observations about the particle deformation in a stable pair:

- The lagging particle is more deformed when it is softer.
- For the same value of La, the lagging particle is more deformed when in a pair than as a single particle.
- Despite having a fixed Laplace number (La_lead_ = 100), the leading particle features a deformation with a weak dependence on the softness of the lagging particle; the leading particle is less deformed when the lagging particle is softer.
- Both particles are more deformed when in a linear pair than when in a staggered pair.

**Fig. 4.**
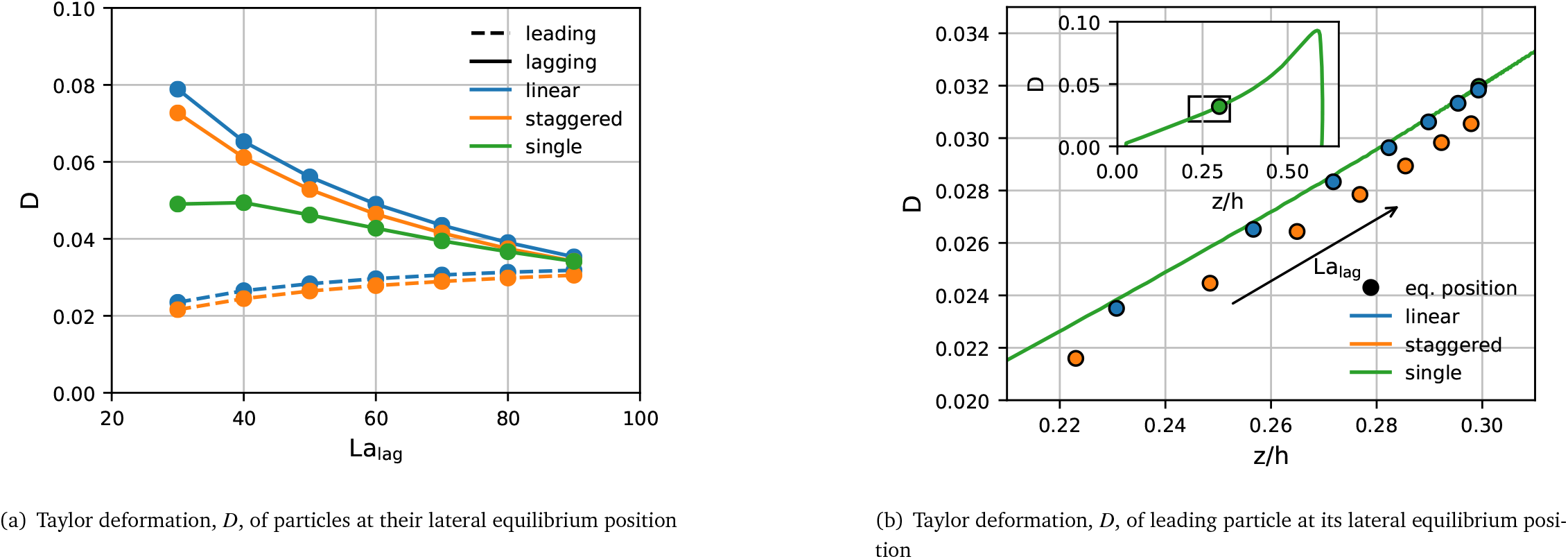
(a) Taylor deformation, *D*, of particles at their lateral equilibrium positions within a stable pair with La_lead_ = 100 and varying La_lag_ *∈* [30, 100]. Leading (dashed line) and lagging (solid line) particles in the linear (blue) and staggered (orange) arrangements are compared with a single particle (green). Note that data is plotted against La_lag_ while La_lead_ = 100 is fixed. (b) Taylor deformation of the leading particle (La_lead_ = 100) for each simulated value of La_lag_ plotted against its lateral equilibrium position *z*_eq_. Both linear (blue) and staggered (orange) arrangements are shown. The green line in the main panel and in the inset shows the reference curve *D*_ref_(*z*) obtained from the transient simulation of a single particle with La = 100 which migrates to its lateral equilibrium position.

While these observations suggest a link between particle deformation and equilibrium behaviour, it is not clear if the change in particle deformation is caused by the particle softness directly or by the shift in the lateral equilibrium position shown in Fig. 3(a), *i.e*., a particle located farther away from the channel centre is expected to be more deformed due to the curvature of the velocity profile and the higher viscous stresses.

To disentangle the effects of lateral position and particle softness on the resulting deformation within a stable pair, we first focus on the leading particle with La_lead_ = 100. We start by establishing a reference curve *D*_ref_(*z*) for the Taylor deformation of a single particle with La = 100. To this end, we perform two separate single-particle simulations with different initial positions along the *z*-axis, one located close to the channel centre and one close to the channel wall. In both simulations, the particle migrates to the same lateral equilibrium position. We assume that the deformation time scale is much shorter than the migration time scale and that the particle deformation is in equilibrium at each point of the trajectory. Thus, the curve in the inset in Fig. 4(b) provides the desired reference *D*_ref_(*z*) at La = 100. The main panel of Fig. 4(b) shows the zoomed-in region of interest of *D*_ref_(*z*) (also indicated by the black box in the inset).

In Fig. 4(b), we compare the actual equilibrium deformation *D* of the leading particle in each of the stable pairs with the reference curve *D*_ref_(*z*). The main observation is that the actual deformation of the leading particle in a linear pair (blue) closely matches *D*_ref_(*z*) for the same lateral position *z*, thus indicating that the lateral position is the main determinant of the particle deformation. However, the leading particle in a staggered pair (orange) is slightly but consistently less deformed than *D*_ref_(*z*) predicts. We can conclude that the deformation of the leading particle is largely predicted by the deformation of a single particle with the same value of La at the same position while the type of pair configuration plays only a minor role. It is beyond the scope of this work to establish how, in a stable pair, the softness of the lagging particle affects the lateral equilibrium position of the leading particle.

#### Transient behaviour

Fig. 5(a) shows the time evolution of the Taylor deformation *D* and the lateral particle position *z* of the leading particle in both the linear and the staggered arrangements for La_lead_ = 100 and La_lag_ = 30. Since the leading particle is initialised without deformation (*D* = 0) at *z*_eq_*/h ≈* 0.3, there is a short time (nearly vertical curve segment) during which the particle adjusts to the local shear stress, reaching a deformation value of *D ≈* 0.032, in both the linear and the staggered arrangements.

**Fig. 5.**
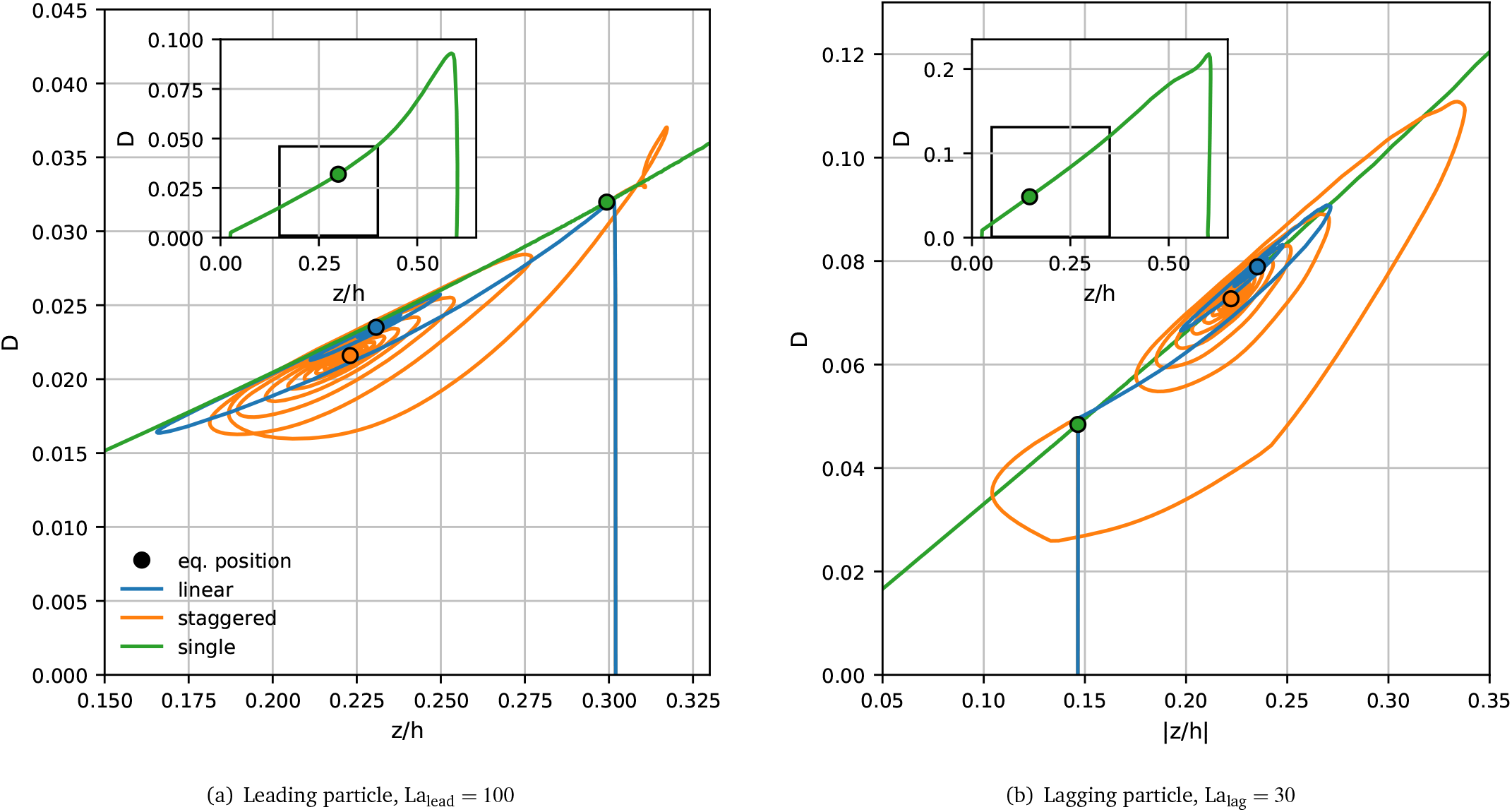
Transient Taylor deformation, *D*, versus lateral position *z* for a pair with La_lead_ = 100 and La_lag_ = 30 in both the linear (blue) and staggered arrangement (orange). (a) Behaviour of the leading particle (La_lead_ = 100) and (b) behaviour of the lagging particle (La_lag_ = 30). The reference curve *D*_ref_(*z*) for a single particle with (a) La = 100 and (b) La = 30 is shown as green line. The main panels show the regions of interest while the insets show the whole reference curves. The leading particle is initialised with *D* = 0 at *z/h ≈* 0.3 and the lagging particle with *D* = 0 at |*z/h*| *≈* 0.15, corresponding to the respective single-particle equilibrium positions.

The subsequent behaviour is different for both arrangements. In the linear pair (blue line), the deformation of the leading particle closely traces the *D*_ref_(*z*) curve (green line) while it oscillates and eventually converges to its equilibrium value (blue circle) close to the reference curve. In contrast, the deformation of the leading particle in the staggered pair (orange line) shows a more pronounced departure from the reference curve with a deviation between *D* and *D*_ref_(*z*) of up to 25%. In equilibrium, the leading particle in the staggered arrangement is slightly closer to the channel centre and the deformation *D* is less than the reference value at the same lateral position (orange circle). In both cases, the leading particle focuses closer to the channel centre than a single particle for the same value of La (green circle). We hypoth-esise that the deformation of the leading particle in the staggered arrangement is more strongly affected by the presence of the lagging particle since the inter-particle spacing is smaller than in the linear arrangement.

Fig. 5(b) shows the transient behaviour of the lagging particle in the same pair as in Fig. 5(a). Here, the green curve is the reference *D*_ref_(*z*) for a particle with La = 30. The initial position of the lagging particle equals the lateral equilibrium position of a single particle with the same value of La, *z*_eq_*/h ≈* 0.15. As in the case of the leading particle, the lagging particle initially adjusts to the local shear stress before oscillating and converging to its equilibrium configuration. It can be seen that the lagging particle in the linear pair (blue line) closely follows the reference curve, just as the leading particle does. The lagging particle in the staggered arrangement (orange line) shows an even stronger deviation from the reference curve (up to 50%) than the leading particle. We conclude that both particles in a forming pair affect each other more strongly in the staggered than in the linear arrangement, probably due to a closer spatial proximity.

### 3.3 Transition between scatter and capture

The transition between pair scatter and capture can be understood in terms of both particle softness and initial particle position.

#### Role of particle softness

We first investigate the transition between pair scatter and capture upon changing La_lag_. The Laplace number of the leading particle is kept at La_lead_ = 100, and particles are initialised at their respective La-dependent single-particle lateral equilibrium position with an axial distance of *δx*_0_ = 10*a*. The trajectories of selected pairs in the linear and staggered arrangement are shown in Fig. 6. Fig. 6(a) depicts the trajectories of the lagging particle in both the linear and staggered arrangement, and Fig. 6(b) and Fig. 6(c) show the trajectories of the leading particle in the linear and the staggered arrangement, respectively. It can be seen that, for both the linear and staggered arrangement, the trajectory type (see Sec. 2.4 for classification) transitions from *Pass and Scatter* to a *Capture* trajectory as La_lag_ increases. Note that, as explained in Section 3.2, the case La_lag_ = La_lead_ = 100 in the linear arrangement does not form a stable pair and is not shown.

**Fig. 6.**
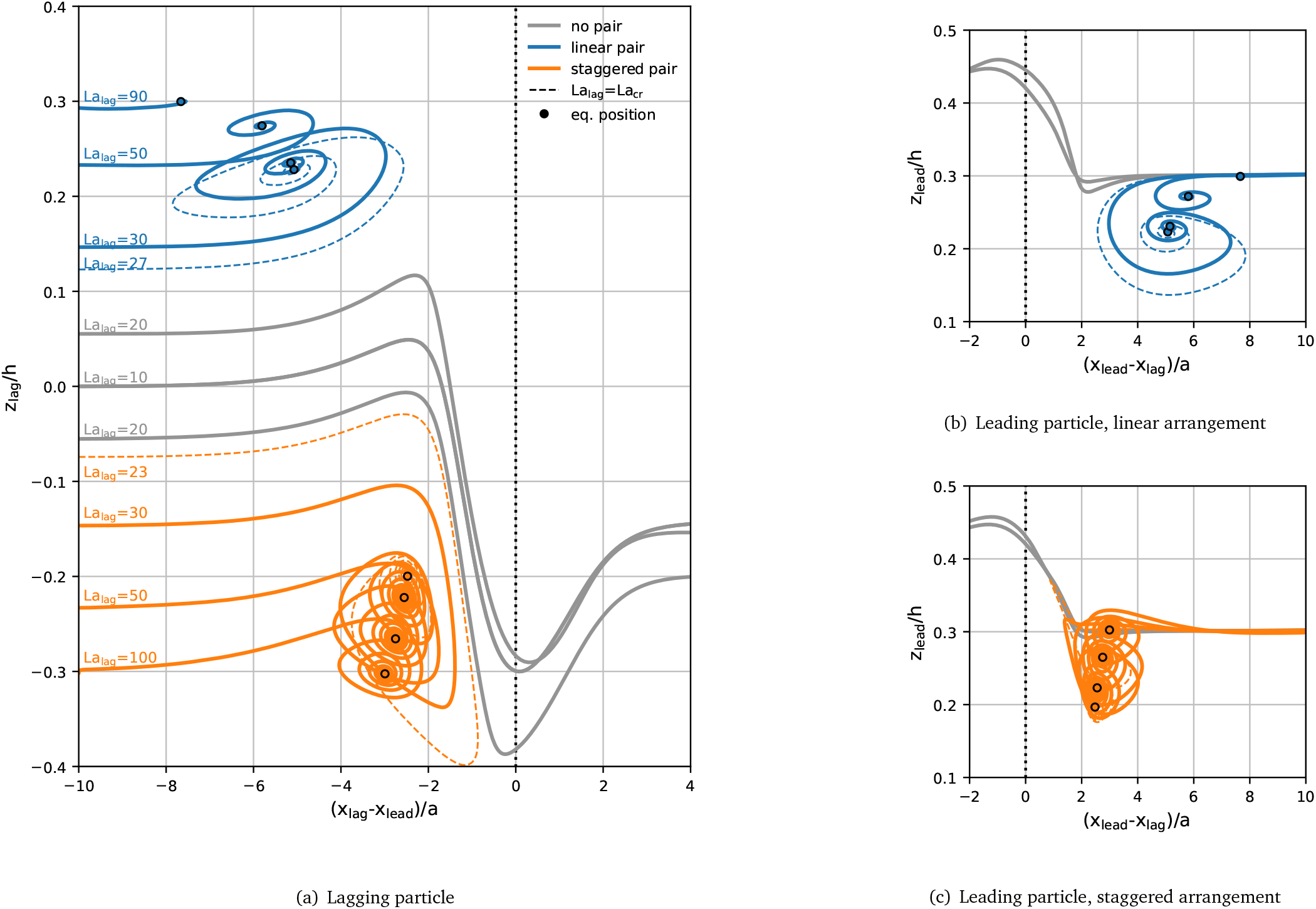
Trajectories of (a) the lagging and (b,c) the leading particle for La_lead_ = 100 and various values of La_lag_. Grey curves denote scatter events, blue curves the formation of a linear pair, and orange curves the formation of a staggered pair. Dashed lines indicate the curves at the critical Laplace number La_cr_ for which a pair just forms. The final equilibrium positions of the particles in a stable pair are indicated by circles. Lines are annotated with the Laplace number of the lagging particle in (a). Particles are initialised at their respective La-dependent single-particle lateral equilibrium positions with an axial distance of *δx*_0_ = 10*a*. For better visibility of the trajectories of the leading particle, the (b) linear and (c) staggered arrangements are shown in separate panels. The vertical black dotted lines indicate the axial position of the leading particle in (a) and the lagging particle in (b) and (c).

All scatter trajectories (grey) lead to the staggered arrangement, irrespective of the initial arrangement, *i*.*e*., two particles in the *Pass and Scatter* mode that are initially on the same side of the channel end up on opposite sides. We do not observe any *Pass and Scatter* events resulting in a linear arrangement. In a denser suspension of particles where multiple encounters between the same two particles might occur, the preference for staggered arrangements might favour the emergence of staggered (rather than linear) trains, a hypothesis that could be tested in the future.

The trajectories of particles that form a captured pair show damped oscillations, both axially and laterally, until a stable equilibrium configuration is reached. These spiral trajectories have been reported in previous studies of particle pairs ^19,36,37^. We observe that, as La_lag_ increases, the oscillation amplitude decreases. This decrease coincides with the increasing equilibrium spacing *δx*_eq_ visible in Fig. 3(b) and in Fig. 6(a,b). Presumably, the oscillations are less pronounced for larger *δx*_eq_ since particles interact less strongly when they are farther away from each other.

The transition between trajectory types occurs between La_lag_ = 20 and La_lag_ = 30 for both arrangements. We define a critical Laplace number of the lagging particle, La_cr_, where the transition from scatter to capture occurs. We find that La_cr_ *≈* 27 in the linear and La_cr_ *≈* 23 in the staggered arrangement. The difference between both critical Laplace numbers corresponds to *≈* 15% variation in softness, well within the natural variation in cell properties ^38^. So far it is unclear whether the transition from scatter to capture is caused by the particle softness directly or by the softness-dependent initial particle position.

#### Role of initial particle positions

The transition point between scatter and capture types coincides with the largest gradient in lateral equilibrium position as shown in Fig. 3. Therefore, the trajectory type might depend on the initial lateral position, irrespective of particle softness. To test this hypothesis, we perform two additional simulations in the *Leading Stiff* configuration. In both simulations, the leading particle remains at La_lead_ = 100, while the lagging particle has either La_lag_ = 20 or 30. Note that the critical Laplace number for both the linear and staggered configuration lies between these two values. We swap the initial positions of the lagging particle in both simulations such that 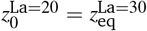 and 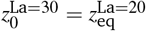. In Fig. 7 we compare these additional trajectories to those obtained when the lagging particle is initially located at its own lateral equilibrium position. We can see that La_lag_ = 20 now leads to a capture trajectory and La_lag_ = 30 to a scatter trajectory.

**Fig. 7.**
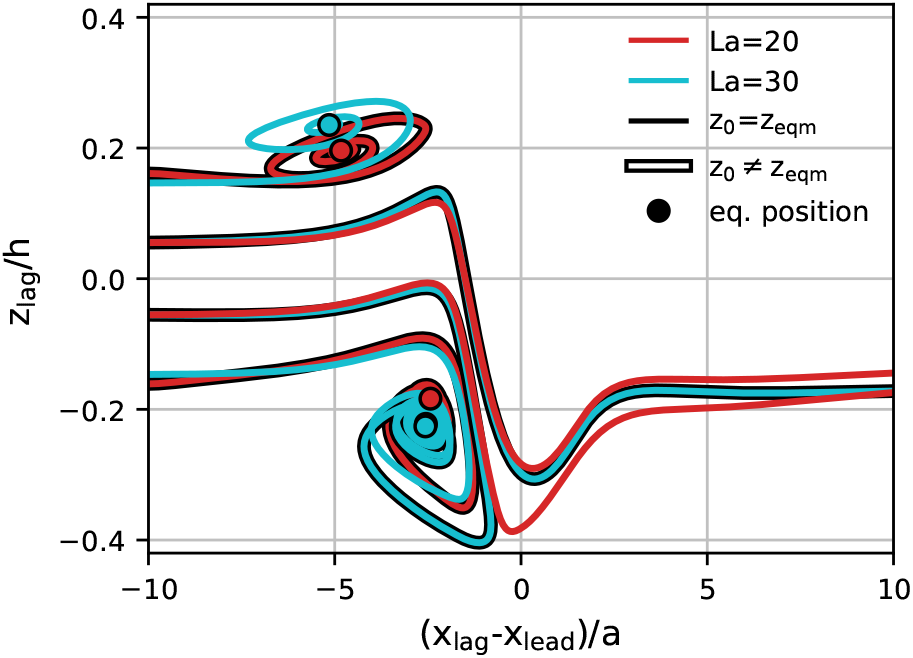
Effect of initial position on pair formation. The leading particle has La_lead_ = 100 (not shown), and the lagging particle has either La_lag_ = 20 (red) or 30 (cyan) in the linear (*z*_0_ *>* 0) and staggered arrangement (*z*_0_ *<* 0). The initial position of the lagging particle is either according to its single-particle equilibrium position or swapped such that 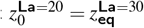 and 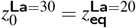.

These results show that the likelihood of a pair forming from particles of different softness also depends on the spatial configurations of the particles at the time of their encounter. In a dilute suspension, particle interactions are rare and it can be assumed that particles have time to equilibrate. If a scatter event happens, particles are likely to migrate back to their equilibrium positions before another particle interaction occurs. However, in a denser suspension, particle interactions occur more often and, therefore, after a scatter event occurs, particles are less likely to have migrated back to their equilibrium positions before interacting with another particle. As a result, the range of spatial configurations at the time of particle interactions is larger in a dense suspension. In our previous work, we have shown that the formation of homogeneous soft particle pairs strongly depends on the spatial configuration during the encounter ^19^. We have also shown that, in the case of heterogeneously sized particles that would not form pairs under equilibrium conditions on approach, stability bands exist where a lagging particle in an off-equilibrium position may form a pair with a leading particle that is in equilibrium ^18^. Thus, we hypothesise that particle softness plays are larger role in pair formation in dilute suspensions while spatial configuration and softness both contribute in denser suspensions.

#### Disentangling the roles of softness and initial positions

To disentangle the effects of initial position and particle softness, we investigate the transient particle deformation for all cases from Fig. 7. Fig. 8 shows the resulting deformation parameter *D versus* instantaneous axial inter-particle spacing *δx* and lateral particle position *z*. We compare the deformation of each particle to the reference deformation *D*_ref_(*z*) of an identical single particle (dotted lines). We make several observations:

**Fig. 8.**
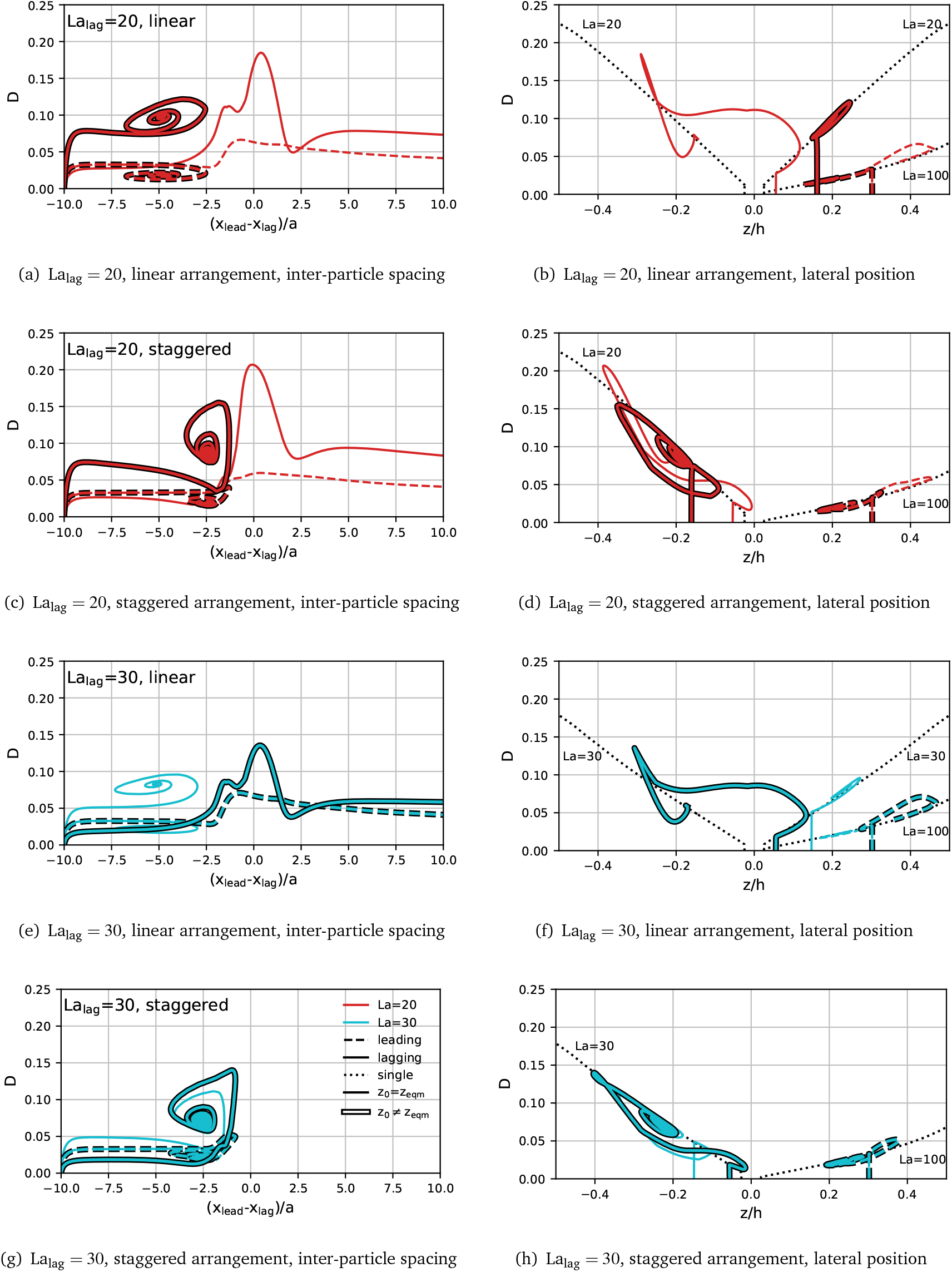
Transient Taylor deformation *D versus* instantaneous axial inter-particle distance *δx* (left column) and lateral particle position *z* (right column). The simulations are identical to those in Fig. 7. The leading particle (dashed line) has La_lead_ = 100 in all cases. The lagging particle (solid line) has either La_lag_ = 20 (red) or 30 (cyan). The black dotted lines show the reference curves *D*_ref_(*z*) obtained from the transient simulation of a single particle with La = 20, 30 and 100 (labelled).

- For the linear arrangement, if a pair forms, both particles closely trace the deformation characteristics *D*_ref_(*z*) of a single particle with the same value of La, independently of the initial position of the lagging particle.
- For the staggered arrangement, if a pair forms, both particles tend to be less deformed than *D*_ref_(*z*) predicts, until both particles form a stable pair, independently of the initial position of the lagging particle.
- For all cases where a pair does not form, the deformation of the particles becomes significantly larger than *D*_ref_(*z*) at some point in time.

We conclude that the actual deformation of the particles plays an important role in the pair formation process, in particle during scatter events.

To demonstrate the role of deformation in pair formation further, we compare the actual deformation with the expected deformation for a given lateral position using the single-particle reference curves *D*_ref_(*z*) for the leading and lagging particles during their interaction. Fig. 9 shows the transient Taylor deformation of the leading and lagging particles with respect to axial interparticle spacing for La_lead_ = 100 and La_lag_ = 20. The actual transient Taylor deformation profiles are identical to those included in Fig. 8(a) and (c). Initial linear and staggered arrangements are considered along with the lagging particle being initialised at its lateral equilibrium position and off-equilibrium. Reference curves *D*_ref_(*z*) for the leading and lagging particles are shown in dotted lines. The grey-shaded area shows the region where the axial centre-to-centre distance between both particles is less than the undeformed particle diameter, thus indicating the region where particle silhouettes partially overlap when observed along the *z*-axis.

**Fig. 9.**
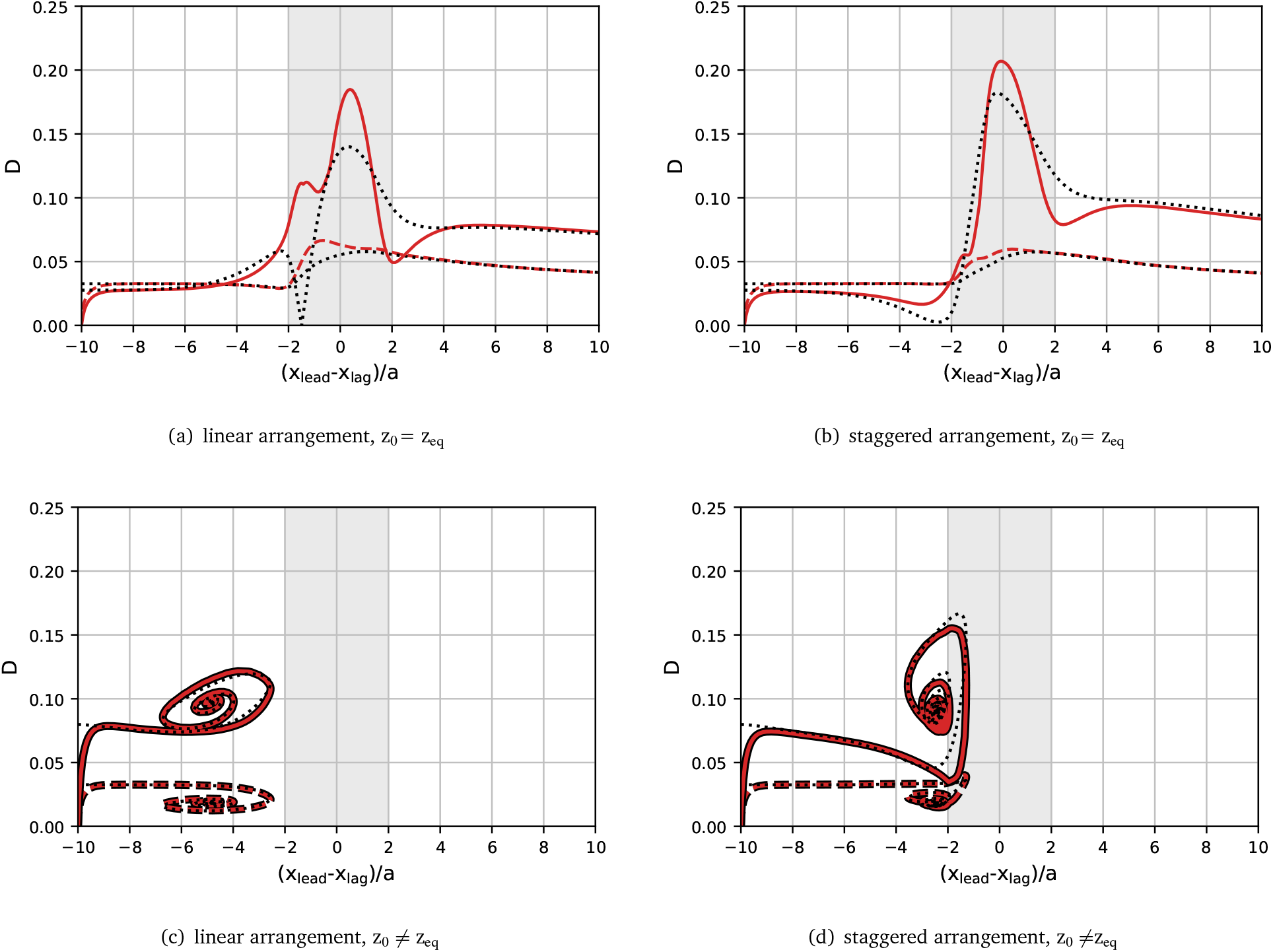
Transient Taylor deformation *D* versus instantaneous axial inter-particle distance *δx* for leading (dashed) and lagging (solid) particles compared to the expected deformation using the reference curves *D*_ref_(*z*) (dotted) obtained from the transient simulation of a single particle. The reference curves for the leading and lagging particles are La=100 and La=20 respectively. The grey area indicates the inter-particle distance where the particles overlap in their undeformed state.

We first consider the situations where no stable pair forms in Fig.9(a) and (b). From Fig. 8, we have established that, when a stable pair does not form, the deformation of the particles becomes larger than predicted by the reference curve *D*_ref_(*z*). Here, we observe that, once |*δx*| *<* 2*a*, the deformation of both particles is larger than predicted by the reference curve. However, as long as |*δx*| *>* 2*a*, the actual deformation profile closely follows that of the reference curve. Once the lagging particle overtakes the leading particle and begins to move away, the actual deformation returns to following the reference curve. From the available data, we cannot conclude whether the scatter events happen because particles are strongly deformed or particles deform more strongly because they overtake each other and need to negotiate the available space.

Finally, we consider the situations where stable pairs form, with initial linear and staggered arrangements in Fig.9(c) and (d), respectively. In the staggered case, we observe that the actual deformation profile follows the reference curve closely, however, some deviations occur during the damped oscillation. This deviation can be attributed to the small axial distance (*δx <* 2*a*) between the particles during the oscillation. In the linear arrangement, the axial distance between stable pairs is around double that of a staggered pair ^35^. As a consequence, the axial distance between the particles does not reach below 2*a* and the actual deformation closely matches the reference curve for the entire trajectory.

### 3.4 Implications for reduced-order models

Our results show that, unless particles are very close (*δx <* 2*a*), their deformation is essentially determined by that of isolated particles with the same properties at the same location. If a pair forms in a linear arrangement, particles in our investigated range do not reach distances smaller than 2*a*, while pairs in a staggered arrangement only briefly reach distances smaller than 2*a* during damped oscillation, before reaching an axial equilibrium distance larger than 2*a*. Although the trajectories of particles can be strongly affected by the presence of other particles over wider distances, the actual particle deformation only deviates from the reference curve derived from the behaviour of a single particle when particles are very close. Our findings, thus, open up the opportunity to develop reduced-order models for particle behaviour in pairs and trains, largely based on the behaviour of a single particle.

Asmolov ^39^ developed a model for the inertial lift force on a rigid, spherical particle:

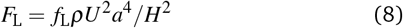

where *f*_L_ is the lift coefficient depending on Reynolds number and the lateral position of the particle. However, the lift coefficient also depends on the particle shape which varies in the case of a deformable particle. Our finding that the particle deformation during pair formation largely follows the behaviour of a single particle could be exploited to modify reduced-order models, such as Eq. (8), to include the effect of transient particle shape on inertial lift even in situations where particles are not isolated.

## 4 Conclusions

The formation of pairs of particles or cells of different types in microfluidic channels can be desired (for example, for the generation of compound particles) or detrimental (for example, for the separation of different particle species). Particle pairs can be classified into staggered pairs (particles located on opposite sides of the channel) and linear pairs (particles located on the same side of the channel). It has been reported that linear particle pairs do not form over a wide range of parameters when both particles have the same properties. However, in reality, cells are generally heterogeneous. Earlier work has shown that a slight heterogeneity in size can lead to the formation of linear pairs. It is still unclear what role the heterogeneity of particle softness plays in the formation of pairs.

We use an in-house lattice-Boltzmann-immersed-boundary-finite-element solver to simulate a pair of particles with different softness flowing through a pressure-driven straight channel with a rectangular cross-section. Particle softness is characterised by the Laplace number, La, where the rigid limit corresponds to La *→* ∞. Both linear and staggered arrangements are considered. We distinguish cases where the leading particle is softer (*Leading Soft*) and where the leading particle is stiffer (*Leading Stiff*). Apart from the outcome of particle interactions, either scattering (no pair forming) or capturing (pair forming), we investigate the transient lateral positions, axial inter-particle spacing, and particle deformation. In order to compare particle behaviour to that of a single particle in isolation, we introduce reference curves consisting of the deformation profile of a single particle as it migrates across the channel cross-section.

We find that softness heterogeneity significantly affects the pair dynamics. Particles generally do not form a pair when the leading particle is softer since the softer particle tends to equilibrate closer to the channel centre where it moves faster and, thus, away from the lagging particle. When the leading particle is more rigid, the outcome of the interaction depends on the softness of the lagging particle: if the lagging particle is sufficiently soft, it catches up with the leading particle and overtakes it, without a pair forming. When the softness of the lagging particle is reduced beyond a critical Laplace number, it still catches up with the leading particle but is able to form a stable pair. For any pair that forms, both particles have nearly identical lateral equilibrium positions, even if their softness is significantly different. Interestingly, the softness of the lagging particle can alter the lateral equilibrium position of the leading particle. Also, we find that the softness of the lagging particle strongly affects the equilibrium axial interparticle spacing in the linear arrangement, while the distance in the staggered arrangement is nearly independent of the softness of the lagging particle.

We find that, for the same value of La, the lagging particle is more deformed when in a pair than as a single particle. Despite having a fixed Laplace number in our study (La_lead_ = 100), the leading particle is less deformed when the lagging particle is softer. It is also observed that both particles are more deformed when in a linear pair than when in a staggered pair. The primary mechanism for the change in particle deformation in a pair compared to an isolated particle is caused by the shift of the lateral equilibrium position to a region with different shear stress. In particular for staggered pairs, the presence of the other particle has an additional non-trivial effect on the deformation of the other particle.

The outcome of the particle interaction depends on both particle softness and initial particle positions. We demonstrate that, when pairs do not form, particles tend to deform more than when they are in isolation. We find that particles ending up in linear pairs closely match the deformation behaviour of a single particle at a given lateral position during pair formation. For staggered pairs, however, the single-particle curves do not predict the particle deformation well since particles approach each other more closely during the formation of a staggered pair.

Our work has important implications for the understanding of collective particle dynamics in inertial microfluidics and the design of microfluidic devices for particle manipulation. For example, our finding that particle deformation follows the behaviour of a single particle at the same location if the other particle is at least one diameter away brings us closer to the development of reduced-order models. Improved reduced-order models are crucial for the accurate simulation of inertial microfluidics in less computationally demanding simulations where particles are not fully resolved.

## Author Contributions

Benjamin Owen: Conceptualization (equal); Formal analysis (lead); Methodology (equal); Validation (lead); Writing – original draft (lead); Writing – review & editing (equal). Krishnaveni Thota: Visualization (supporting); Writing – review & editing (supporting). Timm Krüger: Conceptualization (equal); Formal analysis (supporting); Funding acquisition (lead); Methodology (equal); Project administration (lead); Writing – original draft (supporting); Writing – review & editing (equal).

## Acknowledgements

This work used the Cirrus UK National Tier-2 HPC Service at EPCC (http://www.cirrus.ac.uk). TK received funding from the European Research Council (ERC) under the European Union’s Horizon 2020 research and innovation program (803553).

